# Expression of putative effectors of different *Xylella fastidiosa* subspecies/strains reveals recognition and defense activation in various model plants

**DOI:** 10.1101/2021.05.27.445625

**Authors:** Matthaios Sertedakis, Konstantinos Kotsaridis, Dimitra Tsakiri, Ana Dominguez-Ferreras, Vardis Ntoukakis, Panagiotis F. Sarris

## Abstract

The re-emergence of Gram-negative bacterium *Xylella fastidiosa* in Europe in 2013 impelled the scientific community to discover novel strategies for crop protection. The wide host range of *Xylella* indicates the existence of yet not characterized pathogenic mechanisms to overcome plant defenses. The recent uprising accuracy of a variety of bioinformatics tools, with the ability to predict the function of putative microbial protein represent a useful approach for understanding which of these proteins are associated with pathogens virulence. In this study we collected a number of putative effectors from two *X. fastidiosa* strains: Temecula1 and CoDiRo and the subspecies (ssp.) Sandyi Ann-1. We designed an *in-planta Agrobacterium* based expression system that drives the expressed proteins to the cell apoplast, in order to investigate their ability to activate defense in various model plants. Furthermore, we organized the resulted proteins according to their sequential and structural similarities via the I-TASSER online tool. We identified that various *X. fastidiosa* proteins were able to differentially elicit cell death-like phenotypes in *Nicotiana tabacum, N. sylvestris* and *N. benthamiana*. These proteins are members of different enzymatic groups: a) hydrolases/hydrolases inhibitors, b) serine proteases and c) metal transferases. Collectively, we identified structurally similar proteins that were able to differentially elicit cell death-like phenotypes in different cultivars of the same species. Our findings provide the bases for further studies on the mechanisms that underlie host-defense activation by *X. fastidiosa* putative effectors, as well as, pathogens adaptation in susceptible hosts.

## INTRODUCTION

Plants respond to invading pathogens by exploiting their innate immunity system. microbe-associated molecular patterns (MAMP)-triggered immunity (MTI) and effectors triggered immunity (ETI) have been described as the main two layers of defense during the infection of a host (Jones, J.D.G., 2006; Duxbury et al., 2016; Mermigka et al., 2019; Cui et al., 2015). Recent studies have proposed a revised version of the zig-zag model of plant innate immunity introduced by Jones, J.D.G., 2006 (Jones & Dangl, 2006); strongly indicating the existence of a crosstalk between MTI and ETI. ETI potentiates MTI immune responses and vice versa (Katagiri & Tsuda, 2010; Ngou et al., 2021). MTI is responsible for the detection of pathogen/microbe associated molecular patterns (P/MAMPs) and/or danger associated molecular patterns (DAMPs), via specific cell surface-localized pattern-recognition receptors (PRRs) (Couto & Zipfel, 2016). MTI shields the plant cell from various pathogen-derived molecules or by recognizing self-made elicitors (Malukani et al., 2020) and triggering downstream signaling events to activate defense responses (Schwessinger & Zipfel, 2008; Apama et al., 2009).

MTI defensive outcomes include production of reactive oxygen species (ROS), calcium influx, activation of mitogen-activated protein kinases (MAPKs), chromatin remodeling, differential regulation of gene expression and callose deposition. Collectively, these responses restrict pathogens on the site of the infection and prevent disease development (Lu et al., 2015; Mur et al., 2008; Pardal et al., 2021; Stotz et al., 2014).

In order to reach their nutritional needs and proliferate effectively inside the host, several pathogens have evolved to secrete virulence factors - known as effectors – directly into the host-cell cytoplasm or into the extremely hostile apoplastic space. In several cases, interaction of apoplastic pathogen effectors with plant PRRs has been associated with induction of immune responses and development of Programmed Cell Death (PCD) (van der Burgh & Joosten, 2019). For instance, the apoplastic effectors Avr2 and Avr4 from the fungal pathogen *Cladosporium fulvum* are recognized by the receptor like proteins (RLPs) *Cf-2* and *Cf-4* respectively in *Solanum lycopersicum* and trigger a strong defense response, including PCD (Song et al., 2009; Postma et al., 2016; Ilyas et al., 2015; Kourelis & Van Der Hoorn, 2018). Similarly, the apoplastic effectors Chp-7 and ChpG of *Clavibacter michiganensis*, elicit PCD when they are secreted to the apoplast, but not when they are expressed in the host-cell cytoplasm (Lu et al., 2015). There are additional virulent extracellular effectors related to PCD phenotypes in plants, however the exact mechanisms of innate immunity underlying their perception remain deeply uncharacterized (Nissinen et al., 2009; Lu et al., 2015). *X. fastidiosa* (*Xf*) was first described as the causal agent of “Pierce’s disease” (PD) in grapes and it is an extremely dangerous plant pathogenic bacterium worldwide (Mollenhauer & Hopkins, 1974; Hopkins & Purcell, 2002; Chatterjee et al., 2008). *Xf* is a gram negative, slow growing and strictly aerobic bacterium that has been a subject of interest due to its economic impact. *Xf* has an extremely extended host range which consists of more than 300 plant species (Baldi & La Porta, 2017), including *Nerium oleander, Olea europaea* and *Vitis vinifera* species (Schneider et al., 2020; Food & Authority, 2018; Huang et al., 2020). While there are emerging studies assessing the life style of this pathogen, host specificity and colonization strategies, less progress has been accomplished in the field of the molecular host-pathogen interactions. Similarly, the individual role(s) of putative virulence proteins secreted by *Xf*, in order to subvert host’s immune machinery and how this leads to disease development and finally plant death, is still poorly understood (Roper et al., 2019; Rapicavoli et al., 2018; Chatterjee et al., 2008; Zhang et al., 2015; Nascimento et al., 2016; Gouran et al., 2016).

*Xf* lacks a Type III translocation system (T3SS); the common bacterial transporter of virulence factors from the pathogen’s cytosol directly into the host’s intracellular environment. However, *Xf* possesses Type I, II, IV and V secretion systems (Simpson et al., 2000; Van Sluys et al., 2003). The *Xf* 12-protein Type II secretion system (T2SS) with origins to its close relatives of the *Xanthomonas* group, considerably acts as the main source of its pathogenicity (Rapicavoli et al., 2018). Proteases and cell wall degrading enzymes (CWDEs) are often secreted by T2SS, while mutations on essential components of the secretion mechanism, usually lead to avirulent phenotypes (Rapicavoli et al., 2018).

In this study, using the KEGG database, we searched for homologues of various known type II effector genes of pathogenic microorganisms that could be present in several *Xf* sequenced genomes (**Suppl. Table S1**). This process resulted in selection of nineteen putative *Xf* type II effectors originating from two strains and one spp. for further study (**Suppl. Table S1**).

Gene evolution is a process that involves mechanisms such as gene duplications and horizontal gene transfers, which resulted in the hypothesis that sequence unrelated genes may have high similarity in their tertiary folding and furthermore have the same function in pathogen virulence (de Guillen et al., 2015; Andrie et al., 2008). Based on this hypothesis and using I-TASSER online server, we compared the protein with the highest sequence similarity with the one, which was used as a template for the predicted structures of the selected *Xf* proteins (Roy et al., 2010; Yang et al., 2015; Yang & Zhang, 2015) (**Fig. 1** & **Suppl. Fig. S1**).

**Figure 1.**
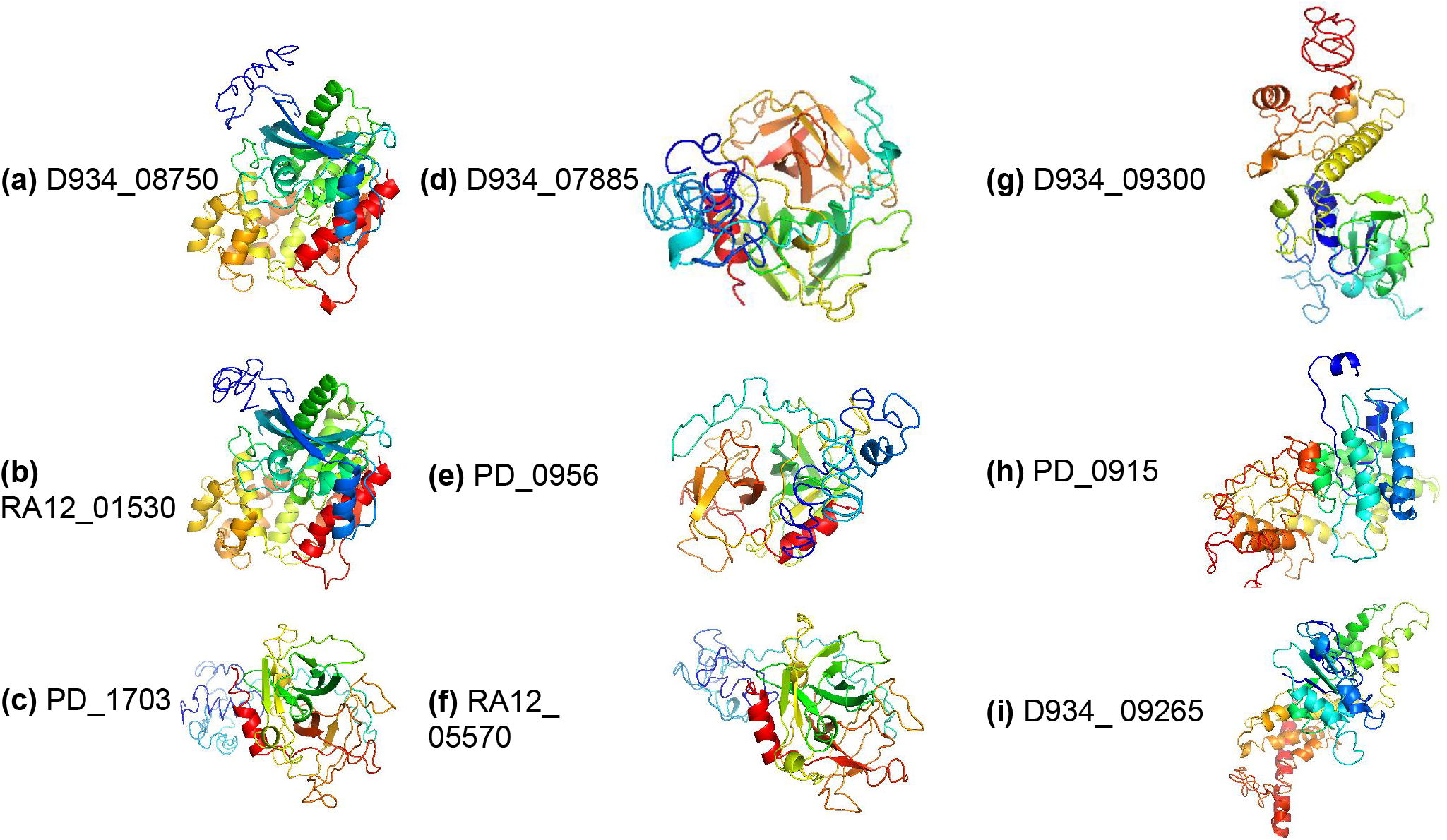
Predicted model presentation of the selected *Xylella fastidiosa* putative effectors using I-TASSER online server. The proteins presented here successfully elicited programed cell death (PCD) in at least one plant cultivar/species tested. The colors suggest the protein orientation (Blue: N-termini, Red: C-termini). **(a-b)** Hydrolase/ Esterase (LipA), **(c-f)** Hydrolase and **(g-i)** Zonular Occludens Toxin, according to their sequence similarities. We used Pymol v2.3.1 to visualize the structures (Schrodinger & DeLano., 2020).

## MATERIALS & METHODS

### Plant material

Three wild type (WT) *Nicotiana* species were used for our study; *N. tabacum* cultivars N34/4, Xanthi, Petit Gerard, *N. sylvestris* ecotypes ITB626, NIC6, A34750352, A04750326, NS25 and TW136 and *Nicotiana benthamiana*. All plants were grown under greenhouse conditions, at 23°C and under a 16-hour photoperiod.

### Bacterial strains

*Escherichia coli* (strains Stellar and DH10b) were routinely grown on LB medium with the appropriate antibiotics and incubation at 37°C for 16 hours (h). Liquid cultures were grown at 37°C for 16 h applying shaking at 200 rpm.

*Agrobacterium tumefaciens* (strains AGL1, C58C1 and GV3101) were grown on LB medium plates with selective antibiotics and incubation at 28°C for two days. Liquid cultures were grown at 28°C for 24h while shaking at 200 rpm.

Available genomic data banks for three strains of *Xylella fastidiosa* (strains Temecula1, CoDiRo and Sandyi ann-1) allowed DNA synthesis of effector genes.

### Cloning & Constructs

All 19 effectors of *Xylella fastidiosa* (originating from Temecula1, CoDiRo, Sandyi ann-1 strains) were synthesized and introduced in plasmid vector pICH41308, kindly provided by Dr. Vardis Ntoukakis (Department of Life Sciences, University of Warwick). Effector genes along with a 35S promoter, fused to the coding sequence of secretion peptide of tobacco PR1a (Pathogenesis-Related protein 1a) and NOS terminator were transferred in binary vector pICH86966 and cloned using Golden Gate cloning (Engler et al., 2008). pMDC:spC7HPB construct, kindly provided by Professor Jane Glazebrook (Department of Plant Biology, University of Minnesota), was introduced in *Agrobacterium tumefaciens* AGL1 and was used in this study as a positive marker of apoplastic HR-like cell for *in planta* assays. pBluescript::PR1a_sp, pICH86988::XopQ:YFP and pICH86988::GUS:YFP plasmids that were used in this work had already been constructed for a previous study in our lab (Michalopoulou et al., 2020).

### *Agrobacterium*-mediated transient expression *in planta*

Transformed *Agrobacterium* cells were grown on LB agar plates with selective antibiotics for 2 days. Single colonies were used to inoculate LB liquid medium containing selective antibiotics and were cultured for 24h. Cells were then harvested by centrifugation, washed with 5 ml of 10mM MgCl_2_ and re-suspended in 1 ml MM solution (10mM MgCl_2_, 10mM MES [pH 5.6]). OD_600_ was adjusted to 0.5 with MM and the bacteria were then used for infiltration of plant leaves with a blunt end syringe. Six week old plants were challenged with Agrobacterium, leading to *in planta* transient expression of *Xf* effectors, as well as positive and negative markers for identification of HR-like Programmed Cell Death (PCD). Infiltrated plants were incubated under greenhouse conditions (23°C, 16-hour photoperiod) and HR-like cell death was assessed 4dpi.

### *In silico* structural predictions of proteins

Structural predictions were made using the I-TASSER online server. Comparisons were performed between the protein with the highest sequence similarity with the one used as a template for the predicted structures of the selected *Xf* proteins (Roy et al., 2010; Yang et al., 2015; Yang & Zhang, 2015). The predicted structure with TM-score >5 was considered reliable. Visualization of the structures was carried out through Pymol v2.3.1 (Schrodinger & DeLano., 2020).

## RESULTS & DISCUSSION

To test whether these proteins can elicit PCD after their delivery into the plant apoplast, we first synthesized the corresponding genes including silent mutations where needed to domesticate the sequences making them compatible with the Golden Gate system. Then, we cloned the synthesized genes of interest in an *Agrobacterium*-mediated transient expression system (**Fig. 2a**). The gene expression in this system was under the transcriptional regulation of the constitutive CaMV 35S promoter. To ensure secretion of the protein into the apoplast we fused the selected *Xf* proteins to the secretion peptide of tobacco PR1a (Pathogenesis-Related protein 1a) (Lu et al., 2015). The PR1a secretion peptide is cleaved upon secretion to the apoplast (Lu et al., 2015). We generated 19 such constructs to screen for PCD symptoms by the selected *Xf* proteins. For our screening we used three distinct *Nicotiana tabacum* cultivars (N34’4, Petit Gerard, Xanthi); the *N. benthamiana* and six *N. sylvestris* ecotypes. As a positive control we used the *Clavibacter michiganensis* apoplastic effector Chp7 that has been shown to elicit an PCD response upon its secretion to the apoplast by the PR1a secretion peptide (Lu et al., 2015) (**Fig. 2b** & **Suppl. Fig. S2**). Furthermore, the type III effector XopQ of *Xanthomonas campestris* pv. *vesicatoria*, served as a secondary positive control, due to its known ability to elicit Hypersensitive Response (HR) in both *N. tabacum* and *N. benthamiana* (Adlung et al., 2016). *Agrobacterium-mediated* transient expression of the GUS reporter gene in these species did not elicit cell death and it was used as a negative control for our transient expression assays.

**Figure 2.**
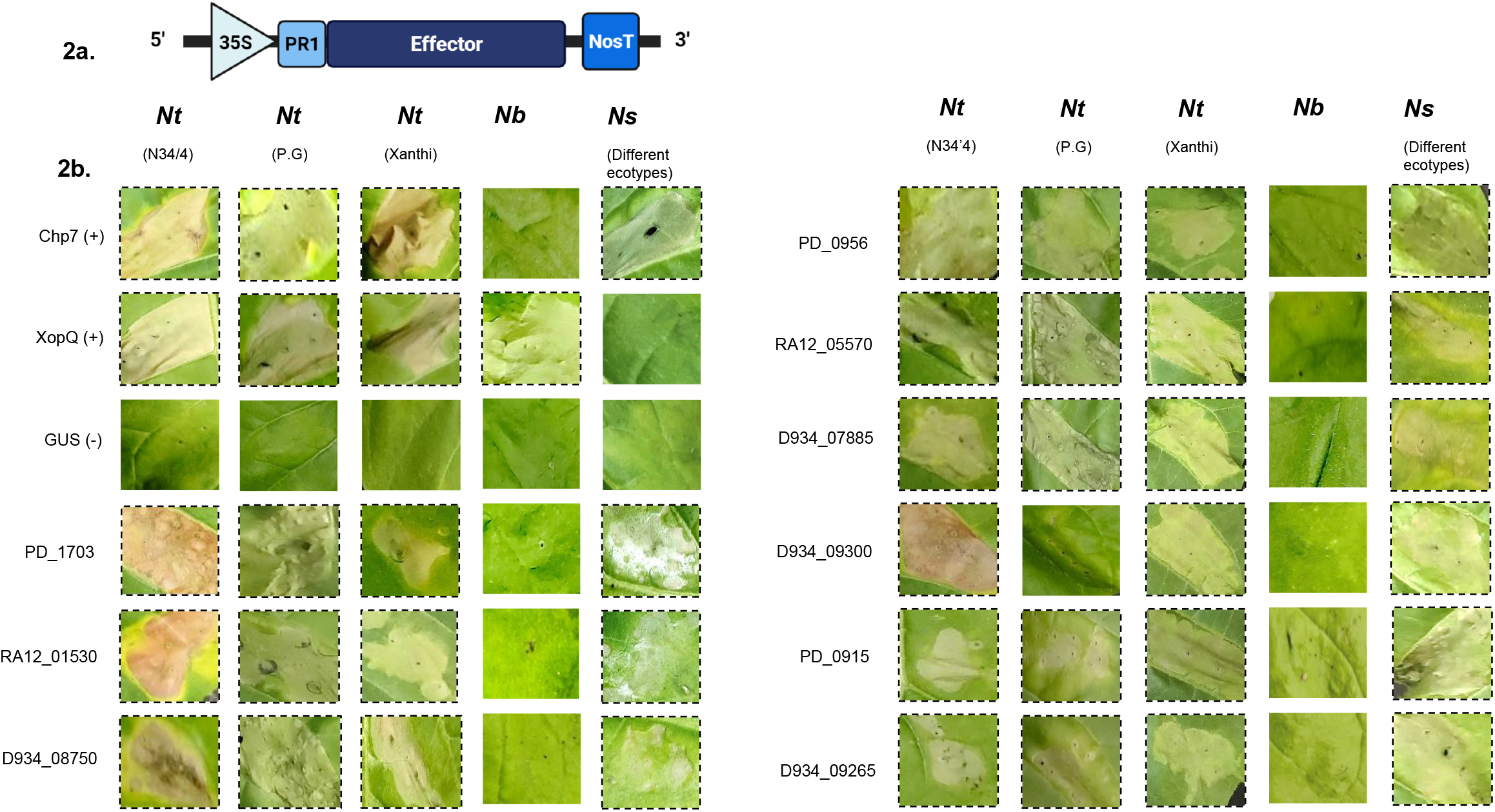
Putative *Xylella fastidiosa* apoplastic effector proteins elicit programed cell death (PCD) in *Nicotiana spp*. **(a)** Schematic representation of the cassette that was cloned in an *Agrobacterium*-mediated plant transient expression system. Genes of interest were under the control of the constitutive CaMV 35S promoter. Secretion of the protein into the leaf apoplast was achieved through fusion to the secretion peptide of tobacco PR1a, which is cleaved during protein secretion (Lu et al., 2015). This figure was created with BioRender.com. **(b)** Apoplastic effector PR1 sp-Chp7 from *Clavibacter michiganensis* along with intracellular acting effector XopQ from *Xanthomonas campestris* pv. *vesicatoria* were used as positive Programmed Cell Death (PCD) - HR markers, while GUS gene served as the negative control for these assays (Lu et al., 2015; Adlung et al., 2016). *X. fastidiosa* virulence factors which induced PCD, following overexpression in the plant apoplast, are illustrated above. PCD occurred 4 days post infiltration under room temperature, in all studied cases. The assays were repeated at least five times for each putative effector with similar results.

Our screening revealed nine proteins that are known or predicted to be type II-secreted by *X. fastidiosa* and were able to elicit PCD phenotypes in different *Nicotiana* species (**Fig. 2b**). Most of these proteins successfully elicited PCD four days’ post inoculation (4 dpi) in all three *N. tabacum* cultivars used and four out of six *N. sylvestris* ecotypes. Interestingly, all nine effectors tested for induction of PCD displayed divergence in distinct *N. sylvestris* ecotypes, while the protein encoded by D934_09300 was able to elicit cell death in *N. tabacum* cv. “N34’4” and cv. “Xanthi” leaves but not in *N. tabacum* cv. “Petit Gerard”, indicating a form of specificity in this response. Moreover, signs of cell death were entirely missing from *N. benthamiana* leaves suggesting that the responses observed in *N. sylvestris* and *N. tabacum* are most likely not a result of cytotoxic effects. Apart from these nine putative effectors, ten more proteins were studied but did not elicit PCD in any plant species/cultivar tested in this study (**Suppl. Fig. S1**).

Cell death phenotypes varied in severity and seemed to develop at different rates. In order to comprehensively evaluate our results, we first assigned a cell death score to each observed cell death phenotype, based on its intensity (**Suppl. Fig. S2**).

We also reviewed the frequency of a certain score regarding both the studied protein and the plant cultivar used in each experiment (**Suppl. Fig. S3**). The necrotic phenotype observed in *N. sylvestris* and *N. tabacum* cv. “N34’4” plants was typically more pronounced compared to that of tobacco cultivars “Petit Gerard” and “Xanthi”. For this, we have to consider potential differences in the transformation efficiency of distinct cultivars under the transient expression system we applied. However, these results could also indicate that the same protein may elicit PCD of varying intensity when introduced to different *Nicotiana* relatives or cultivars of the same plant species and hint that the potency of this type of response is possibly host-dependent.

The *in silico* structural prediction presented here, indicates that all three proteins encoded by PD_0956, RA12_05570 and D934_07885, which successfully elicited PCD in all studied tobacco cultivars and in *N. sylvestris* ecotype A_34750352, have a high structural similarity with hydrolases (**Fig.1**; **Table 1**). Hydrolases form a big distinct enzyme class that includes enzymes which act as biochemical catalysts by using water molecules to break chemical bonds. This class contains enzymes classified as: a) lipases; b) phosphatases; c) glycosidases; d) peptidases; and e) nucleosidases. Specifically, serine proteases/endopeptidases/hydrolases are enzymes where the nucleophilic serine residue in their active center is used for the hydrolysis of their substrates (Simon & Cravatt, 2010). Hydrolases group includes proteases that are secreted by various pathogens having a wide range of functions in virulence. They also constitute an important group of *X. fastidiosa* including cell wall-degrading enzymes (CWDEs) (Nascimento et al., 2016). CWDEs presence in the apoplast can trigger immunity responses, mostly through a modified “self” recognition of degradation products of these enzymes by plant PRRs (van der Burgh & Joosten, 2019). Similarly, serine proteases delivered by pathogens into the apoplast have been shown to activate PCD (Lu et al., 2015). Provided that PD_0956, RA12_05570 and D934_07885 proteins’ putative enzymatic activity is valid, their ability to elicit PCD could be considered a DAMP-recognition event. Serine proteases are also present in large families of plant extracellular proteins that are often involved in signaling pathways associated with pathogen resistance (Hou et al., 2019). Therefore, manipulation of such pathways by bacterial proteases could be a virulence strategy.

**Table 1.** Sequence similar and structural template proteins for the predicted structures of the nineteen selected *Xylella fastidiosa* putative proteins using I-TASSER.

| *Xylella fastidiosa*<br>Proteins | Sequence<br>similarity (PDB) | Description | Structural<br>template (PDB) | Description |
| --- | --- | --- | --- | --- |
| PD_0956 | 3WY8 | Hydrolase/Protease | 3WY8 | Hydrolase/Serine protease |
| PD_0915 | 2R2A | Zonular Occludens Toxin<br>(Zot) | 2DHR | ATP-dependent<br>metalloprotease/Hydrolase |
| PD_1703 | 3WY8 | Hydrolase/serine protease | 1Z8G | Hydrolase/Hydrolase Inhibitor |
| D934_08750 | 3H2K | Hydrolase/Esterase (LipA) | 3H2K | Hydrolase/ Esterase (LipA) |
| D934_07885 | 3WY8 | Hydrolase/Serine protease | 1Z8G | Hydrolase/Hydrolase Inhibitor |
| D934_09300 | 2R2A | Zonular Occludens Toxin<br>(Zot) | 2R2A | Zonular Occludens Toxin ( Zot) |
| D934_09265 | 2R2A | Zonular Occludens Toxin<br>(Zot) | 4WWO | Transferase/Transferase Inhibitor |
| RA12_01530 | 3H2K | Hydrolase/ Esterase (LipA) | 3H2K | Hydrolase/ Esterase (LipA) |
| RA12_05570 | 3WY8 | HydrolaseSerine protease | 1Z8G | Hydrolase/Hydrolase Inhibitor |
| D934_00810 | 5N8P | Membrane Protein | 3JAV | Transport Protein |
| D934_05685 | 5N8P | Membrane Protein | 5IJO | Transport Protein |
| D934_12725 | 3JAV | Transport Protein | 3JAV | Transport Protein |
| D934_08755 | 3H2K | Hydrolase/ Esterase (LipA) | 3H2K | Hydrolase/ Esterase (LipA) |
| D934_12535 | 3H2K | Hydrolase/ Esterase (LipA) | 3H2K | Hydrolase/ Esterase (LipA) |
| D934_12795 | 7KVE | Blood Clotting | 1RWR | Cell Adhesion |
| RA12_11155 | 4UIC | Sugar Binding Protein | 1G6O | Hydrolase |
| RA12_11125 | 3V05 | Toxin | 5N8P | Membrane Protein |
| RA12_03930 | 6VDP | Oxidoreductase | 6VDP | Oxidoreductase |
| RA12_03905 | 6W1S | Gene Regulation | 6WIS | Gene Regulation |

Another prominent group of *Xf* proteins is that of PD_1703, RA12_01530 and D934_08750 that all trigger PCD in three tobacco cultivars and two *N. sylvestris* ecotypes TW_136, NS_25 but not to ITB_626 (**Fig. 2b**) (Zhang et al., 2015). According to our structural analysis (**Table 1**), the last two proteins revealed similarities to LipA, a known *Xanthomonas oryzae* pv. *oryzae* cell wall degrading enzyme (CWDE). While, for the PD_1703, even if it was previously characterized as a LipA-like protein (Nascimento et al., 2016), according to our I-TASSER structural prediction, it revealed similarities to Hydrolase/Serine proteases (**Table 1**). However, hydrolases class is one of the largest and most diverse enzyme families which among others includes proteases and lipases, so, this might be a misannotation of the particular protein database.

LipA homologues are present in all sequenced Xanthomonads and are predicted lipases, although LipA actually exhibits esterase activity (Apama et al., 2009) (**Fig. 3**). LipA is known to elicit immune responses in rice and recent findings point to the possible involvement of a rice Wall-Associated Kinase (WAK) *OsWAKL21.2* in LipA recognition (Jha et al., 2007; Malukani et al., 2020). Structural similarities of LipA with PD_1703, RA12_01530 and D934_08750, could mean that these proteins act and are recognized in a similar manner (this finding is under further investigation by our group). Notably, PD_1703 has been shown to elicit PCD in grapevine a known *Xf* ‘Temecula-1’ host. However, PD_1703 was found to be vital for *Xf* virulence in grapevines, suggesting that other virulence components of the pathogen could potentially suppress the PCD induction (Nascimento et al., 2016).

**Figure 3.**
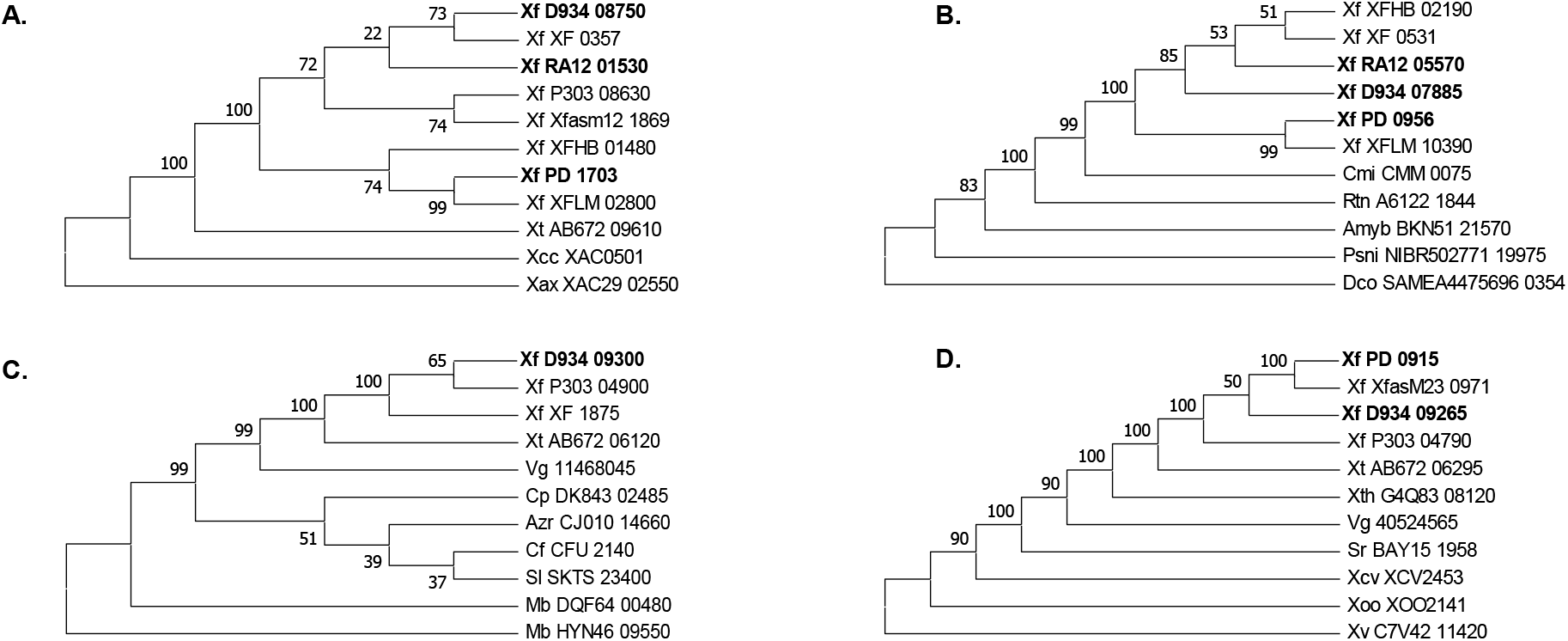
Phylogenetic trees were constructed for all nineteen putative *Xf* effector proteins that are presented in this study, which were divided into subgroups based on their ability to elicit programed cell death (PCD) and on their orthology, according to KEGG database: (**A-D**) Proteins that induced PCD in this study, with a predicted orthology of (**A**) lipases, (**B**) peptidases, (**C-D**) Zona Occludens Toxins. These proteins were correlated with 35 close protein relatives from *Xanthomonas, Clavibacter, Ralstonia, Amycolatopsis, Pseudarthrobacter, Dermatophilus, Streptomyces, Stenotrophomonas, Moraxela, Azoarcus, Collimonas, Sulfurimicrobium, Chromobacterium* genera and viruses *Stenotrophomonas* phage phiSHP2, *Stenotrophomonas* phage SMA6. The evolutionary history in each group presented here was inferred using the Neighbor-Joining method (Saitou & Nei, 1987). The bootstrap consensus tree inferred from 1500 replicates is taken to represent the evolutionary history of the different taxa belonging amino-acid sequences as mentioned before. The evolutionary distances were computed using the Poisson correction method (Zuckerkandl & Pauling, 1965) and are in the units of the number of amino acid substitutions per site. Evolutionary analysis was conducted in MEGA X (Kumar et al., 2018). The abbreviations of microbes and the gene loci used for the construction of these phylogenetic trees are presented in **Supplementary Table S2**.

In this study we also focus on three *Xf* proteins encoded by PD_0915, D934_09265 and D934_09300 that were found to elicit apoplastic PCD in tobacco and one *N. sylvestris* ecotype (**Fig. 2b**; **Table 1**). Structural analysis revealed that this group consists of proteins with sequence and structural similarity to *“Zonula occludens”* toxins or “Zot proteins”, although PD_0915 was predicted to be more confidently similar to a metal-transferase. The “Zot” protein was described first in *Vibrio cholera*, where it is involved in intestinal barrier disturbance, however, “Zot” proteins were identified later in several other pathogens (Pérez-Reytor et al., 2018, 2020) (**Fig. 3**). Zot proteins have been associated with high cytotoxicity before (Pérez-Reytor et al., 2018), though this is not always the case. For instance, in *Vibrio parahaemolyticus*, “Zot” expression did not positively correlate with cytotoxicity, rather than with an actin disturbance on infected cells (Pérez-Reytor et al., 2020). *Xf* putative “Zot” proteins studied here appear not to correlate to cytotoxic effects. Interestingly, the protein encoded by D934_09300 did elicit PCD in the apoplast of *N. tabacum* cv. “N34’4” and cv. “Xanthi”, and *N. sylvestris* ecotype NS_25 but this kind of response was not observed in *N. tabacum* cv. “Petit Gerard”; or in *N. sylvestris* ecotype ITB_626 and in *N. benthamiana* (**Fig. 2b**). These data suggest specific recognition of D934_09300 and highlight the complexity of the plant surveillance system and its possible differentiation among distinct cultivars of the same species.

Finally, ten putative *Xf* effectors, which were unable to induce necrosis during *in planta* assays in the selected hosts, were also analyzed for their tertiary structures using I-TASSER online server and certain predictions could be made for their folding and function (**Suppl. Fig. S1**; **Table 1**). Notably, LipA-like proteins D934_08755 or D934_12535, despite their strong correlation with other cell death inducers described in this study (PD_1703, D934_08750, RA12_01530), were incapable of causing similar phenotypes when expressed in the apoplast of *Nicotiana* species. This potentially indicates putative alterations on their active sites that prevent their binding to specific substrates of plant cell wall.

All the nineteen proteins were used for phylogenetic analysis using homolog proteins obtained from the KEGG database (**Fig. 3**; **Suppl. Fig. S4**).

Our data, collectively, pinpoint nine proteins belonging to the sparsely studied *X. fastidiosa* putative “effectorome” that can elicit PCD when transiently expressed and secreted into the leaf apoplast of different *Nicotiana* species. These proteins are structurally predicted as putative “CWDEs” or “Zot toxins” that originate from different *X. fastidiosa* strains/subspecies. Lack of signs of cytotoxicity, along with the predicted enzymatic activity of these proteins, hints their possible recognition by the plant innate immunity system. At least in one case, the protein eliciting the response is a known required virulence factor of the pathogen, suggesting that it employs other virulence strategies to suppress immune responses and avoid recognition. The suppression of immune responses through type III-delivered effector proteins is a common feature among other members of the *Xanthomonadaceae* family (Jha et al., 2007). However, since *X. fastidiosa* lacks such a system (Rapicavoli et al., 2018), how this bacterium avoids recognition by the host’s surveillance system, remains to be elucidated. In summary, our findings, open possibilities and encourage further investigation and identification of the related PRRs that could be a potential biotechnological tool to confer broad-spectrum disease resistance against *X. fastidiosa*. Quite recently, the expression of a PRR receptor in sweet orange has been shown to confer ligand-dependent activation of defense responses against a citrus infecting strain of *X. fastidiosa* (Mitre et al., 2021).

## Acknowledgements

The project was partially supported and D.T. was supported by the “The Vineyard Roads” (project code: 2018ΣE01300000) part of the “Emblematic Research Action of National Scope for the exploitation of new technologies in the Agri-food sector, specializing in genomic technologies and pilot application in the value chains of “olive”, “grapevine”, “honey” and “livestock”)» financed by Greek national funds through the Public Investments Program (PIP) of General Secretariat for Research & Technology (GSRT) Duration 2019-2021.

K.K. was supported by the General Secretariat of Research and Innovation (GSRI) through the project “Innovative plant protection technologies for quarantine pathogens of *Xanthomonadaceae* family utilizing tools of Optoacoustic and Molecular Biology - INNOVA-PROTECT”, Project code: T1EΔK-01878.

## Authors’ contributions

P.F.S. designed the research. M.S., K.K, and D.T. performed the research. V.D. and P.F.S. analysed the data. V.D. and A.D.F. provided technical support; lab material and tools. M.S., K.K, and D.T. and P.F.S. wrote the paper. All authors have read and approved the manuscript.

## SUPPLEMENTARY DATA LEGENDS

**Supplementary Table S1.** All 19 effector proteins tested for induction of PCD in *Nicotiana* species owe their origins to two pathogenic strains of *Xylella fastidiosa* Temecula1, CoDiRO and ssp. *Sandyi*. Each strain has been associated with infectious diseases in susceptible hosts *Vitis vinifera*, Olive trees and *N. oleander*.

| *Xf* gene ID | Subspecies/ Strains | Host |
| --- | --- | --- |
| PD0915 | strain Temecula1 | Grapevine |
| PD0956 | strain Temecula1 | Grapevine |
| PD1703 | strain Temecula1 | Grapevine |
| RA12_01530 | strain CoDiRO | Olive tree |
| RA12_03905 | strain CoDiRO | Olive tree |
| RA12_03930 | strain CoDiRO | Olive tree |
| RA12_05570 | strain CoDiRO | Olive tree |
| RA12_11125 | strain CoDiRO | Olive tree |
| RA12_11155 | strain CoDiRO | Olive tree |
| D934_00810 | subsp. sandyi Ann-1 | Oleander |
| D934_05685 | subsp. sandyi Ann-1 | Oleander |
| D934_07885 | subsp. sandyi Ann-1 | Oleander |
| D934_08750 | subsp. sandyi Ann-1 | Oleander |
| D934_08755 | subsp. sandyi Ann-1 | Oleander |
| D934_09265 | subsp. sandyi Ann-1 | Oleander |
| D934_09300 | subsp. sandyi Ann-1 | Oleander |
| D934_12535 | subsp. sandyi Ann-1 | Oleander |
| D934_12725 | subsp. sandyi Ann-1 | Oleander |
| D934_12795 | subsp. sandyi Ann-1 | Oleander |

**Supplementary Table S2.** Bacterial species and gene loci members used for phylogenetic analysis presented in **Figure 3** and **Supplementary Figure S4**.

**Supplementary Figure S1.**
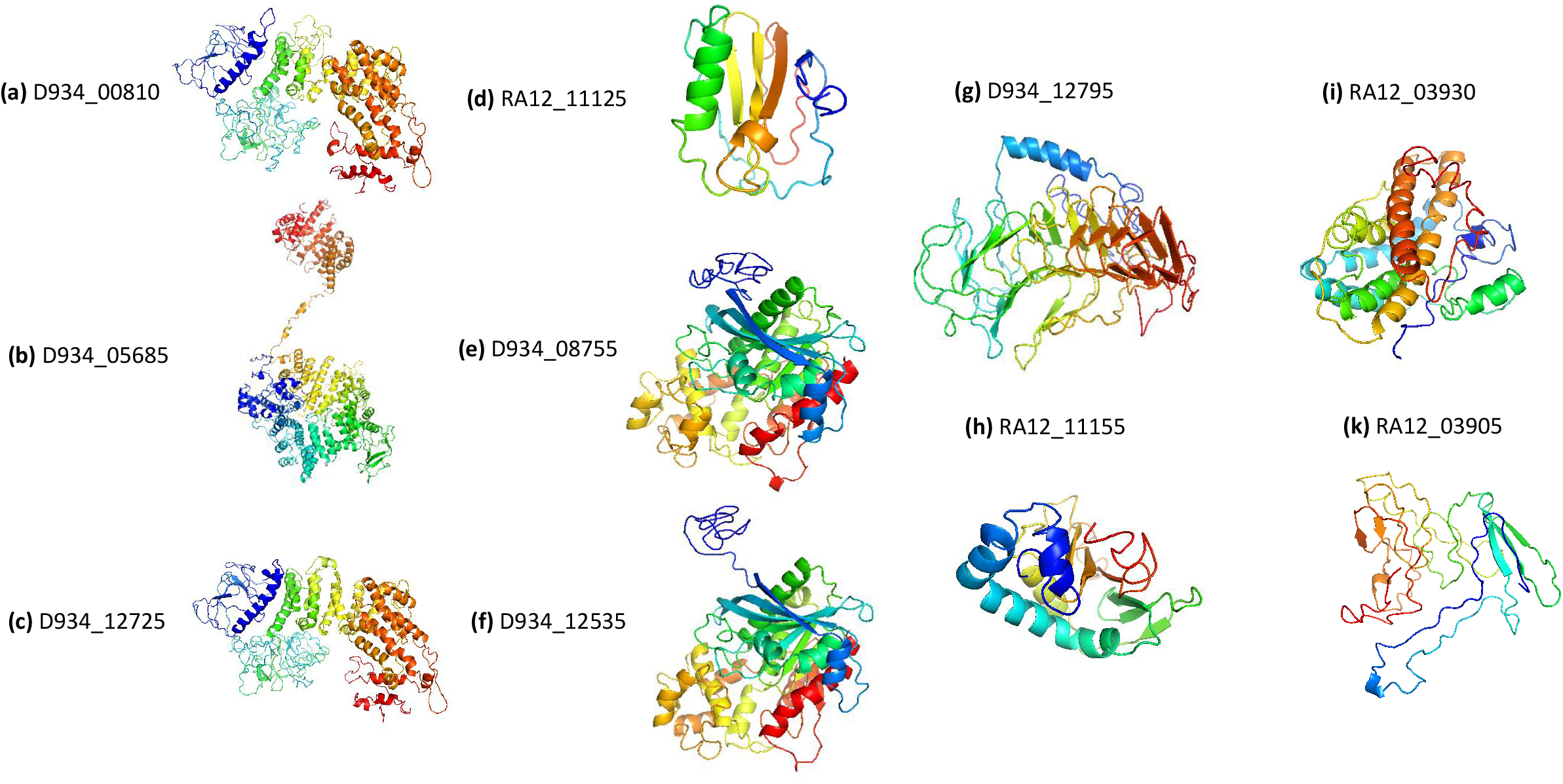
Predicted model presentation of the selected *Xylella fastidiosa* effectors using I-TASSER online server. The proteins presented here did not elicit programed cell death (PCD) in any of the plant cultivar/species tested. The colors suggest the protein orientation (Blue: N-termini, Red: C-termini). **(a-c)** Transport Protein, **(d)** Membrane Protein, **(e-f)** Hydrolase/ Esterase (LipA), **(g)** Cell Adhesion protein, **(h)** Hydrolase, **(i)** Oxidoreductase and **(k)** Gene Regulation. We used Pymol v2.3.1 to visualize the structures. The presented proteins did not induce PCD phenotype in any tested plant cultivars (Schrodinger & DeLano., 2020).

**Supplementary Figure S2.**
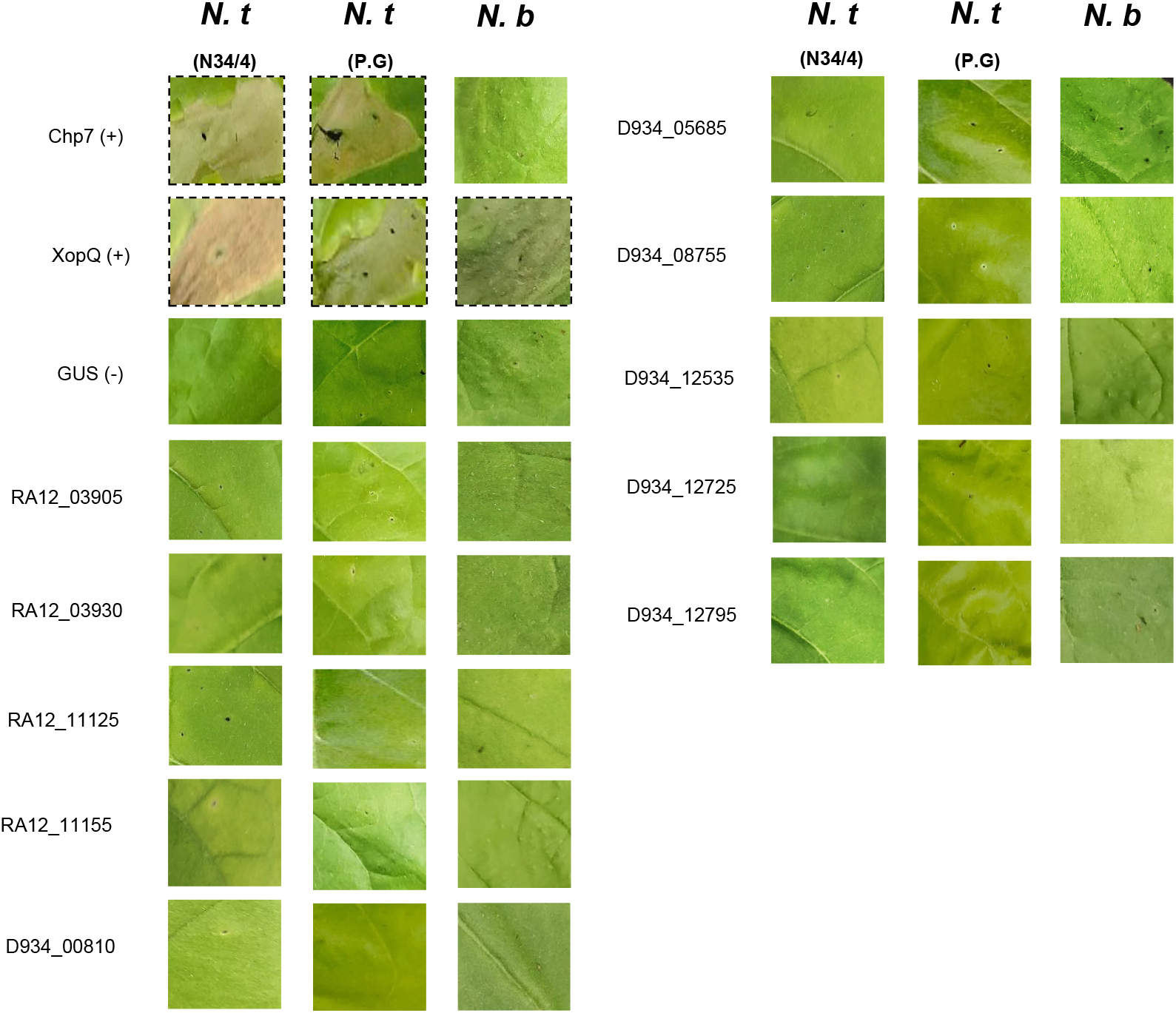
*Xylella fastidiosa* virulence factors which did not elicit programed cell death (PCD), following overexpression in the plant apoplast, are illustrated above. Photographs of infiltrated leaves were taken 4 days post infiltration. In all cases, plants were incubated at room temperature. Apoplastic effector PR1 sp-Chp7 from *Clavibacter michiganensis* along with intracellular acting effector XopQ from *Xanthomonas campestris* pv. *vesicatoria* were used as positive Programmed Cell Death (PCD) - HR markers, while GUS gene served as the negative control for these assays (Lu et al., 2015; Adlung et al., 2016). The assays were repeated at least five times for each putative effector with similar results.

**Supplementary Figure S3.**
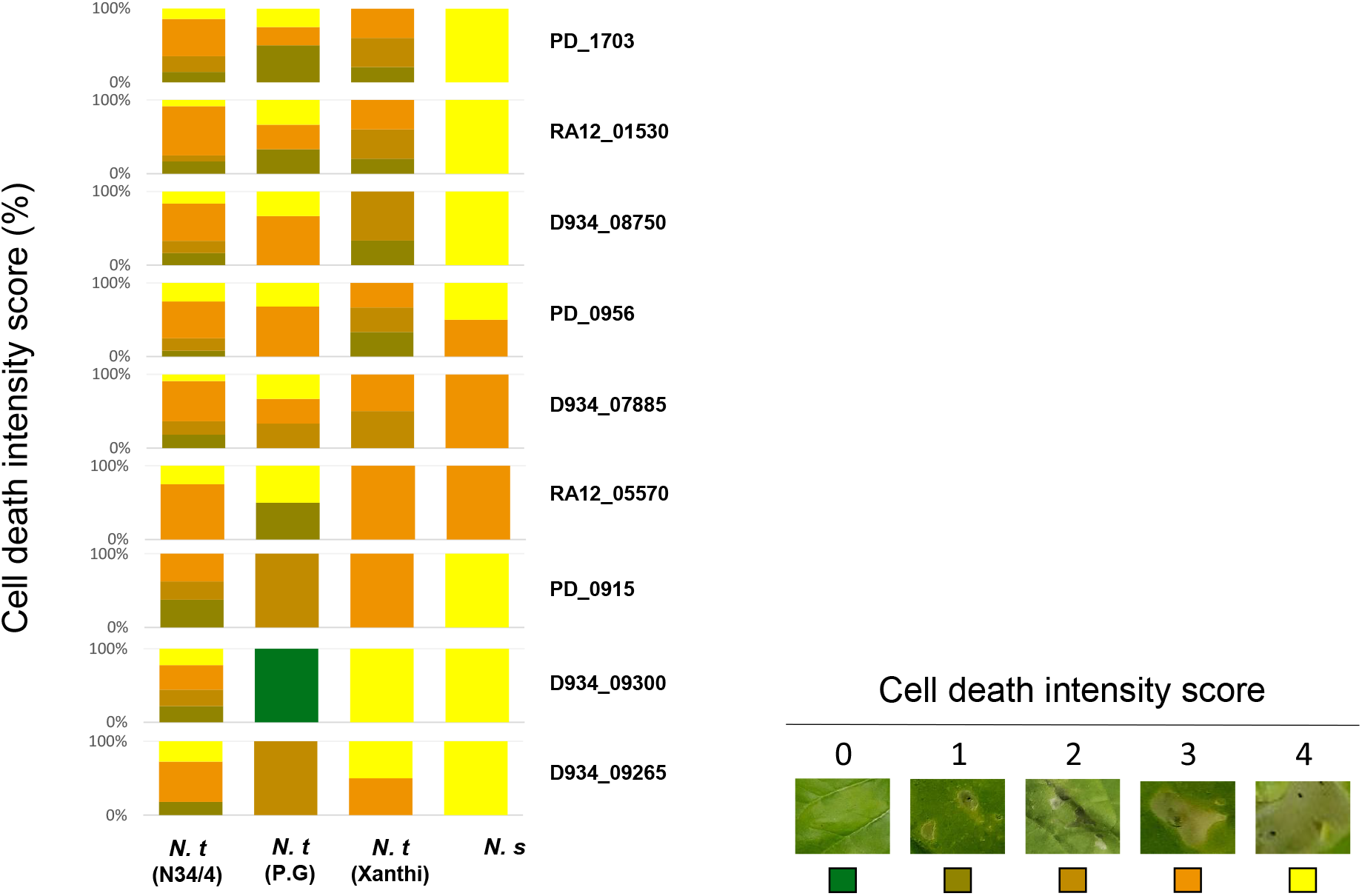
Cell death intensity score. Cell death score percentage (%) in this diagram is representative of all consistent experimental replicates previously introduced in Figure 2b. Diagram bars are color-coded based on a cell death intensity scale 0-4. Colors in each bar represent the cell death score in percentage out of the total infiltrated panels scored for the nine *Xylella fastidiosa* proteins that elicited programed cell death (PCD) in *Nicotiana spp*.

**Supplementary Figure S4.**
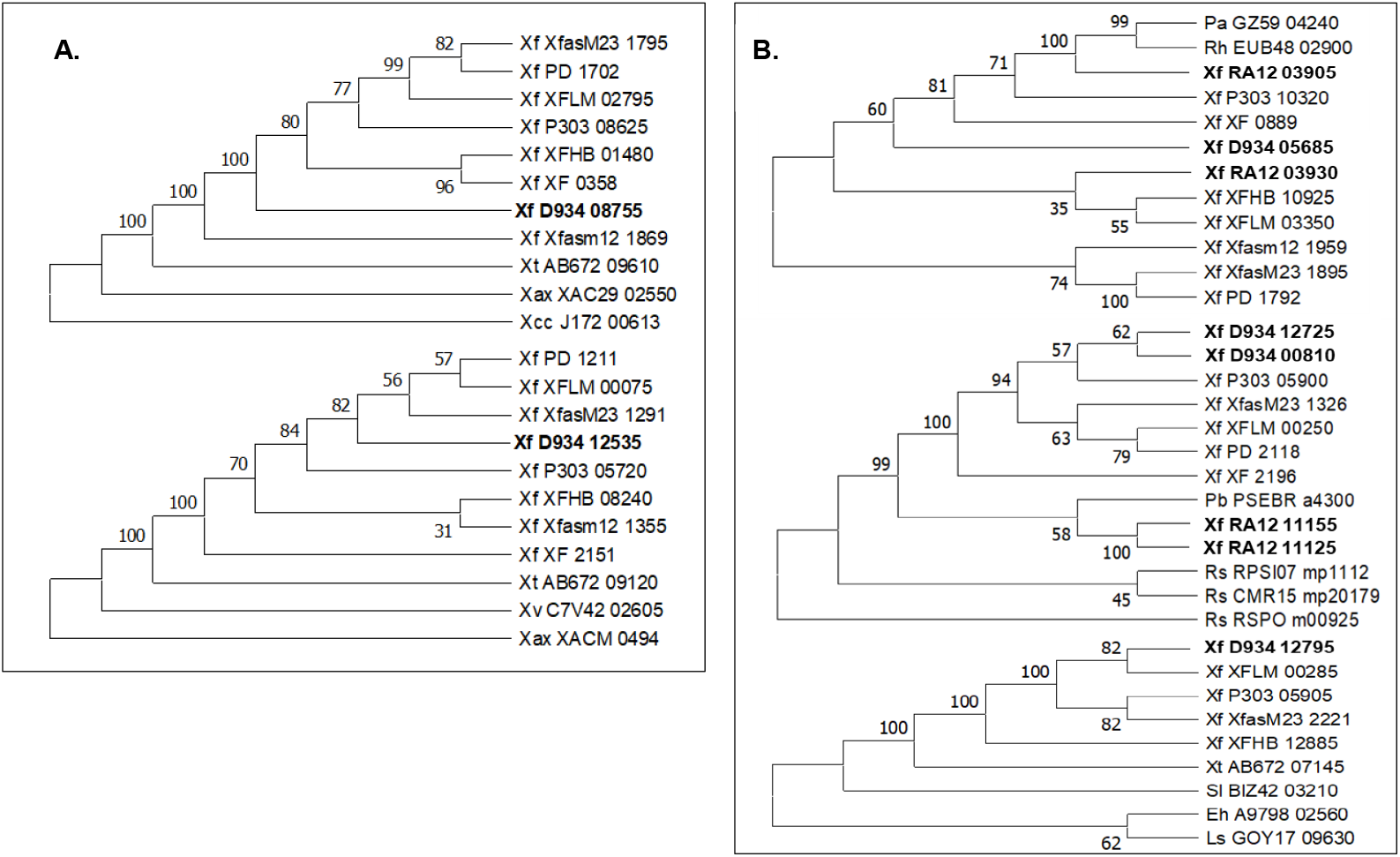
(**A-B**) Proteins that did not induce programed cell death (PCD) in this study, with a predicted orthology of (**A**) lipases and (**B**) haemagglutinins. These groups of proteins from *Xylella fastidiosa* were correlated with 56 proteins with origins to bacterial genera of *Pectobacterium*, *Rhodoferax*, *Ralstonia*, *Pseudomonas*, *Stenotrophomonas, Edwardsiella* and *Lysobacter* (Saitou & Nei, 1987). The evolutionary history in each group presented here was inferred using the NeighborJoining method (Saitou & Nei, 1987). The bootstrap consensus tree inferred from 1500 replicates is taken to represent the evolutionary history of the different taxa belonging amino-acid sequences as mentioned before. The evolutionary distances were computed using the Poisson correction method (Zuckerkandl & Pauling, 1965) and are in the units of the number of amino acid substitutions per site. Evolutionary analysis was conducted in MEGA X (Kumar et al., 2018). The abbreviations of microbes and the gene loci used for the construction of these phylogenetic trees are presented in **Supplementary Table S2.**

